# Glycosylation differentially affects immune cell-specific tetraspanins CD37 and CD53

**DOI:** 10.1101/2023.03.29.534715

**Authors:** Sjoerd van Deventer, Ilse A. Hoogvliet, Merel van de Voort, Frank Arnold, Annemiek B. van Spriel

**Affiliations:** Department of Medical BioSciences, Radboud University Medical Center, Nijmegen, The Netherlands; Department of Cell, Developmental & Cancer Biology, Oregon Health & Science University, Portland, OR, USA

## Abstract

Tetraspanin proteins play an important role in many cellular processes as they are key organizers of different receptors on the plasma membrane. Most tetraspanins are highly glycosylated at their large extracellular loop, but the function of this post-translational modification remains largely unstudied. In this study we investigated the effects of glycosylation of CD37 and CD53, two tetraspanins important for cellular and humoral immunity. Broad and cell-specific repertoires of N-glycosylated CD37 and CD53 were observed in human B cells. We generated different glycosylation mutants of CD37 and CD53 and analyzed their localization, nanoscale organization and partner protein interaction capacity. Abrogation of glycosylation in CD37 revealed the importance of this modification for CD37 surface expression, whereas neither surface expression nor nanoscale organization of CD53 was affected by its glycosylation. CD37 interaction with its known partner proteins, CD20 and IL-6Rα, was not affected by glycosylation, other than via its changed subcellular localization. Surprisingly, glycosylation was found to inhibit the interaction between CD53 and its partner proteins CD45 and CD20. Together, our data show that tetraspanin glycosylation affects their function in immune cells, which adds another layer of regulation to tetraspanin-mediated membrane organization.

## Introduction

Many important cellular processes at the level of the plasma membrane of both healthy and malignant cells are modulated by tetraspanin proteins [1–4]. This family of small four-transmembrane proteins are key membrane organizers that form tetraspanin nanodomains by interacting laterally amongst themselves and with partner proteins [3,5,6]. These interactions are predominantly mediated by the large extracellular loop (also called EC2) in tetraspanins [7,8]. The EC2 of most tetraspanins is equipped with one or more glycosylation groups attached to asparagine residues (N-glycosylation). The highly conserved nature of these N-glycosylation sites and the large size of this post-translational modification, which often composes more than half of the molecular weight (MW) of the protein, suggests that glycosylation may be important for the interactions and function of tetraspanins.

Glycosylation of CD63, for example, has been shown to mediate its interaction with chemokine receptor CXCR4 leading to enhanced trafficking to endosomes and lysosomes and thus downregulation of surface expressed CXCR4 [9]. In addition, CD63 glycosylation controls surface expression of MDR1 (Multidrug resistance protein 1) and has been implicated in breast cancer cell malignancy [10]. On the other hand, glycosylation of CD82 was reported to be important for its interaction with integrin α5β1, and crucial for the metastasis inhibiting potential of CD82 in several cancer models [11,12]. CD82 glycosylation also plays an important role in homing of acute myeloid leukemia to the bone marrow by affecting the nanoscale organization of both tetraspanin CD82 as well as its partner protein N-cadherin [13]. However, the function of glycosylation of tetraspanins in the immune system remains largely unexplored.

In this study we investigated the glycosylation of tetraspanins CD37 and CD53, which are exclusively expressed on immune cells. The expression of CD37 is highest in B cells, where it enables α4β1 integrin-AKT signaling required for long-lived plasma cell survival [14] and is directly involved in the transduction of survival and apoptotic signals [15]. In the malignant counterparts of the B cell, B cell lymphomas, CD37 expression was identified as novel independent prognostic factor [16,17]. This was related to increased IL-6 signaling and fatty acid metabolism in CD37-deficient lymphomas [18,19]. CD37 can be N-glycosylated on three conserved residues in its EC2, the mass of the sugar groups adding up to about 50% of the total molecular weight [20].

CD53 expression is highest in B cells and myeloid cells, but also T cells show a significant expression [21]. CD53 deficiency in both B and T cells results in reduced homing of these cells towards the lymph nodes as a result of L-selectin loss on their cell surface [22]. In B cells, CD53 promotes B cell receptor signaling by recruiting protein kinase C to the plasma membrane [23]. T cell receptor signaling can also be enhanced by CD53 as it functions as a regulator of the highly abundant phosphatase CD45 [24]. CD53 can be glycosylated on two conserved asparagine residues in its EC2 domain and glycosylation represents about half of the total molecular weight of CD53, in line with CD37 [25].

Here we show that CD37 and CD53 contain a broad and cell-specific repertoire of glycosylation, which differentially affects their function. Whereas glycosylation has a clear effect on the localization of CD37, it does not affect localization or nanoscale organization of CD53. We observed glycosylation of CD53 to inhibit its interaction with partner proteins, in contrast to previous studies on CD63 and CD82.

## Results

### Broad and cell-specific repertoires of glycosylated CD37 and CD53

N-glycosylation of CD53 can occur on two highly conserved asparagine residues (N129 and N148) flanking the B helix in its large extracellular loop (Fig 1A, S1A). We investigated glycosylation of CD53 in a panel of different B cell lines and detected a broad band pattern between 25 and 60 kDa on a Western blot (Fig 1B). This band pattern was reduced to a single non-glycosylated band of 20 kDa when N-glycosylation was removed by PNGase-F. All B cell lines harbored a broad repertoire of differentially glycosylated CD53, wherein sugar increases its molecular weight by 25 to 200%. Interestingly, this broad repertoire was cell-specific as clear differences were observed in the CD53 band pattern of different cell lines (Fig 1B, S1B). A range of differentially glycosylated CD53 proteins was also observed in primary B cells isolated from human blood, although the highest MW species (above 40 kDa) appeared to be absent (Fig 1C). CD37 contains three N-glycosylation sites (N170, N183 and N188) close to its variable D-helix that are highly conserved (Fig 1D, S1C). Glycosylation of CD37 appeared as a broad band pattern between 30 and 70 kDa, which reduced to a single non-glycosylated band of 25 kDa upon PNGase-F treatment (Fig 1E). In line with CD53, the CD37 band pattern showed clear differences between different B cell lines and was more narrow for primary B cells isolated from human blood (Fig 1E,F S1D).These data demonstrate the presence of broad and cell-specific repertoires of glycosylated CD53 and CD37 in human B cell lines and primary B cells.

**Figure 1:**
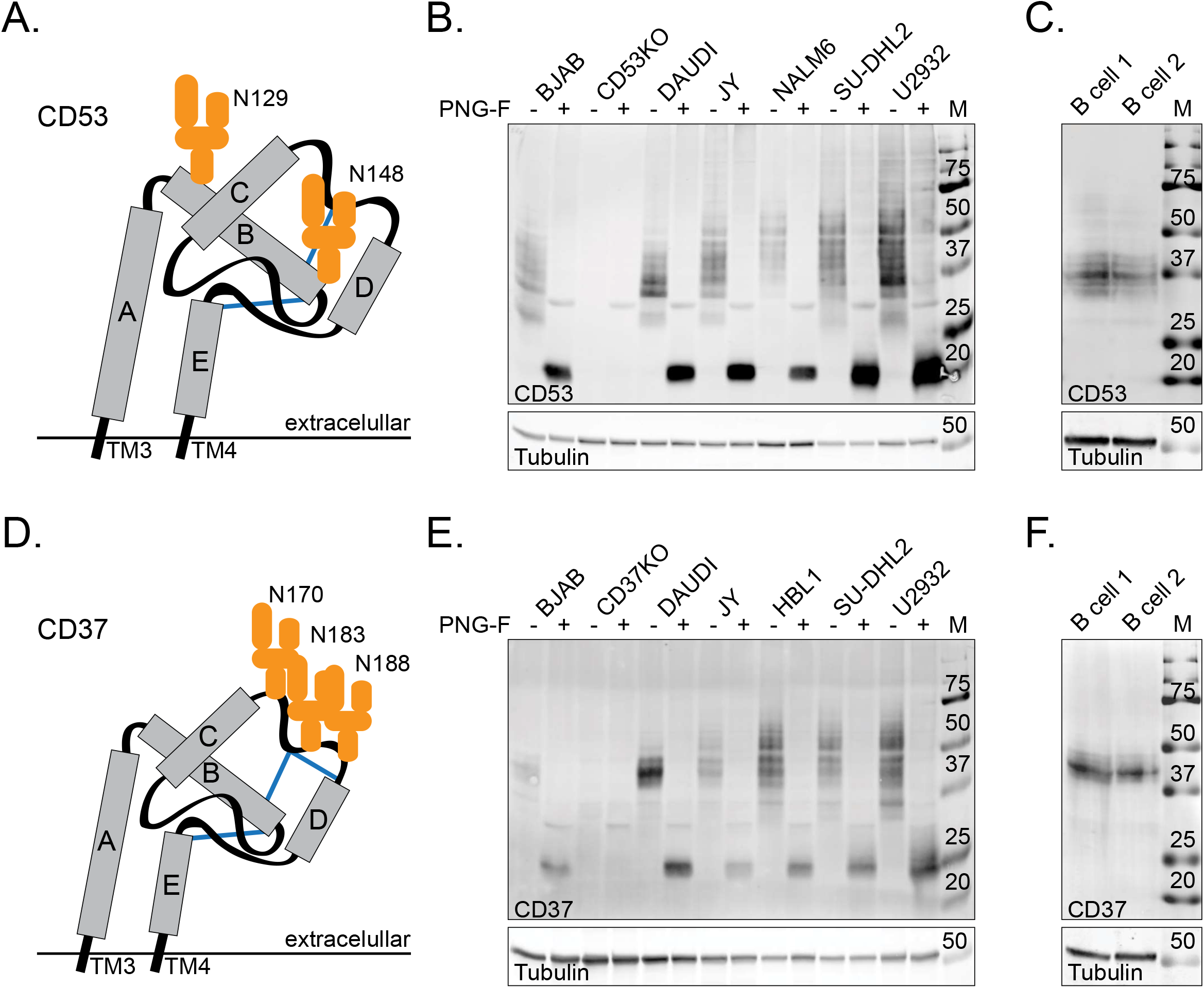
Broad and cell specific repertoires of CD53 and CD37 in B cell lines and primary B cells. Schematic representation of the large extracellular loops of tetraspanin CD53 (**A**) and CD37 (**D**) with 5 alpha helices (A-E) and cysteine bridges (blue lines). Asparagine residues that can be N-glycosylated are indicated. Glycosylation of CD53 (**B**) and CD37 (**E**) is visible on western blot as a broad band pattern (25-70 kDa) in different B cell lines and primary B cells isolated from human blood of healthy donors (**C**,**F**). CD37 and CD53 knock-out (KO) BJAB cells serve as negative control for the staining. Treatment with PNGase-F removes all N-glycosylation leading to a single band around 20 kDa (CD53) and 25 kDa (CD37).

### CD37 surface expression is dependent on its glycosylation

To study the effect of CD37 glycosylation, its three glycosylation sites were mutated (asparagine to glutamine) in a plasmid containing GFP-CD37. Transfection of the non-mutated (WT) parental plasmid in CD37KO BJAB cells resulted in surface expression that was comparable to endogenous levels of CD37 in BJAB cells (Fig 2A). In contrast, the mutated non-glycosylated CD37(NG) was neither detected at the cell surface, nor intracellularly, by a CD37-specific antibody (Fig 2B). To assess whether CD37(NG) was still surface expressed, proteins exposed on the plasma membrane were biotinylated before lysis and immune precipitation (IP) of GFP-tagged CD37 protein. Immunoprecipitated WT GFP-CD37 was observed at the expected size (∽60–75 kDa), but only the higher MW bands (larger glycosylation groups) were biotinylated thus surface exposed (Fig 2C). GFP-CD37(NG), on the other hand, appeared as a sharp band of ∼50 kDa with a weak biotin signal, suggesting low CD37 surface expression (Fig 2C, 2D). Both the low surface expression of CD37(NG) and the over-representation of the higher MW species of CD37 in the biotin signal indicate that CD37 surface expression is affected by its glycosylation. To exclude an effect of the GFP-tag, experiments were repeated with the small ALFA-tag [26] attached to CD37 yielding very similar results (Fig S2A-D). Also, loss of antibody detection of CD37(NG) was independent of the antibody used, as a different CD37 antibody clone yielded the same result (Fig S2E).

**Figure 2:**
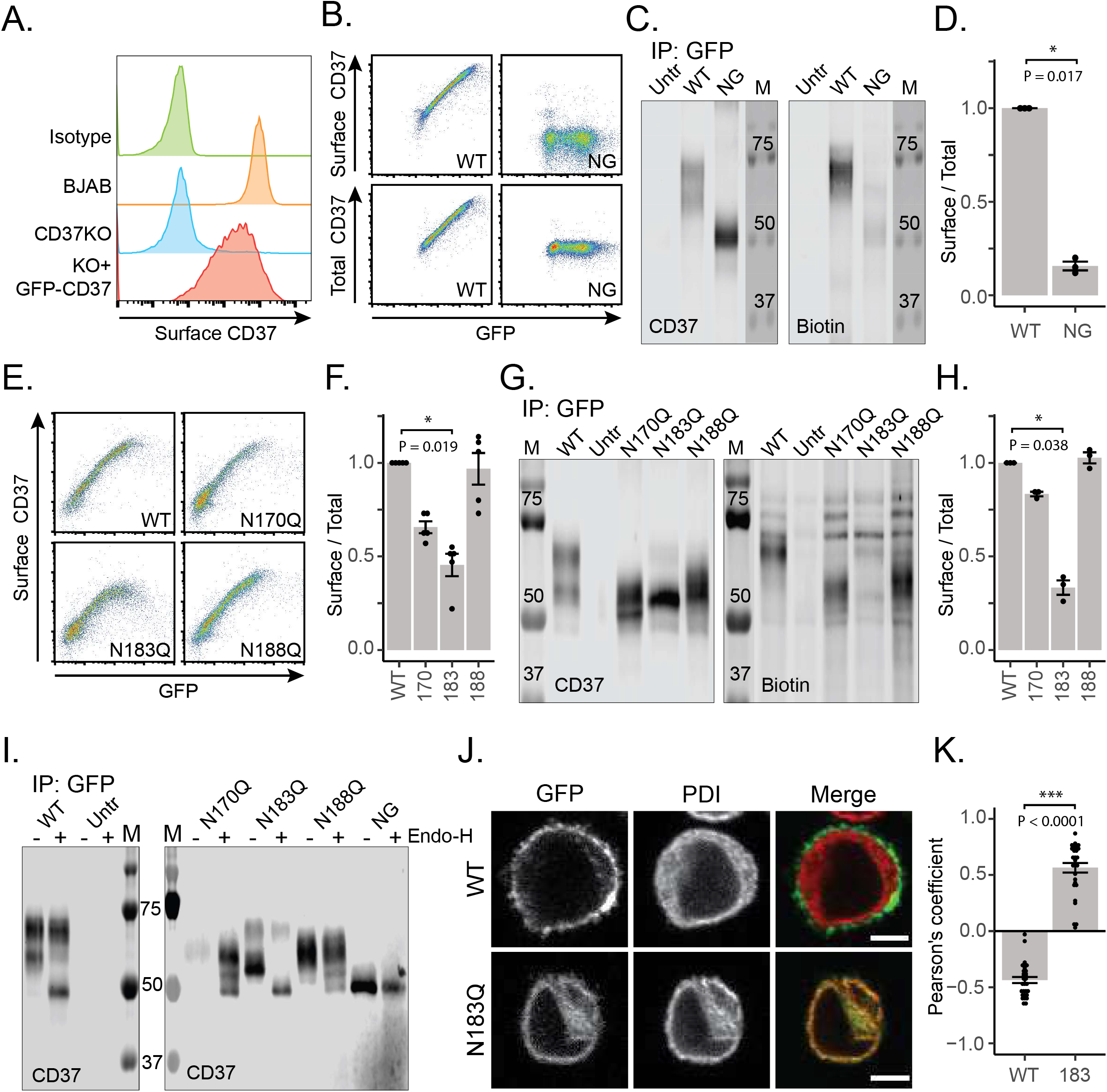
CD37 surface expression is dependent on its glycosylation. (**A**) Endogenous surface expression of CD37 in BJAB cells compared to CD37KO BJAB cells transfected with GFP-CD37 analyzed by flow cytometry. (**B**) Correlation between antibody staining of surface and total CD37 versus GFP signal of CD37KO BJABs transfected with GFP-CD37 (WT) or its non-glycosylated counterpart (NG). (**C**) Immunoprecipitation of WT and NG GFP-CD37 expressed in CD37KO BJAB cells after surface biotinylation. The higher molecular weight (MW) bands of WT CD37 show a strong biotin signal, in contrast to the lower MW bands of WT CD37 and CD37(NG). (**D**) Quantification of (C). Biotin signal (surface) divided by CD37 signal (total) normalized to WT (mean ± sem, non-paired T-test, N=3). (**E**) Antibody staining of surface CD37 versus GFP signal of CD37KO BJAB cells expressing GFP-CD37 (WT) or single-mutants N170Q, N183Q and N188Q. (**F**) Quantification of (E). Antibody signal (surface) divided by GFP signal (total) normalized to WT (mean ± sem, Friedman test and Dunn’s multiple comparison, N=5). (**G**) Western blot of immunoprecipitated WT and single-mutant GFP-CD37 after surface biotinylation and the quantification (**H**) of biotin signal (surface) divided by CD37 signal (total) normalized to WT (mean ± sem, Friedman test and Dunn’s multiple comparison, N=3). (**I**) Endo-H treatment of immunoprecipitated WT, single-mutant and NG GFP-CD37 removes ER-associated, but not Golgi-associated, glycosylation. (**J**) Confocal microscopy showing strong overlap between GFP-CD37(NG) and endoplasmic reticulum marker PDI, in contrast to WT GFP-CD37 and PDI. Scale bar represents 10 μm. (**K**) Quantification of the obtained images showing Pearson’s coefficients of 30 cells per condition (mean ± sem, non-paired T-test, N=3). * P≤0.05 and ***P≤0.001

To reveal whether the glycosylation-dependent surface expression of CD37 could be attributed to a particular glycosylation site, surface expression of the single mutants (N170Q, N183Q, N188Q) was analyzed in CD37KO BJAB cells. We observed a lower surface staining of N170Q and N183Q than expected based on the signal of the attached GFP (Fig 2E, 2F). N188Q, on the other hand, showed a similar ratio between surface staining and GFP signal as the WT. Similar observations were made for ALFA-tagged CD37 single mutants in CD37KO OCI-LY1 cells (Fig S2F, S2G). To exclude that reduced surface recognition of N170Q and N183Q was caused by decreased recognition by the CD37 antibody, CD37KO BJABs expressing WT or single mutant GFP-CD37 were surface biotinylated before lysis and IP via the GFP tag. The biotinylation signal of the single mutants suggested that N183Q is significantly less expressed at the cell surface than WT, whereas N170Q and N188Q showed a similar surface expression (Fig 2G, 2H) in line with the flow cytometry data. The MW of the band pattern of the single mutants was in between the fully glycosylated version of the WT (∼75 kDa) and the MW of non-glycosylated GFP-CD37 (∼50 kDa), suggesting that all three glycosylation sites contribute to the full glycosylation of CD37. All in all, these data show that glycosylation of N183 is mainly required for the surface expression of CD37.

Since the total protein level of N183Q was similar to WT whereas its surface expression was impaired, the mutated CD37 protein has to accumulate in an intracellular compartment. To address whether N183Q was able to leave the ER after synthesis, we subjected immunoprecipitated protein to Endo-H, an enzyme that can cleave ER-associated, but not Golgi-associated, glycosylation. When immunoprecipitated WT GFP-CD37 was subjected to this enzyme, the higher MW proteins (∼75 kDa) were unaffected, whereas the lower MW proteins (∼60 kDa) moved to the non-glycosylated size (∼50 kDa) (Fig 2I). This suggests that the lower MW proteins were localized in the ER, probably on their way to further processing in the Golgi and the plasma membrane. For the single mutants, the band-pattern of N170Q and N188Q were hardly affected by Endo-H, whereas the majority of the N183Q bands shifted to the non-glycosylated size, supporting that this mutant accumulates in the ER. We verified this observation by demonstrating a strong co-localization of N183Q with ER-marker PDI, that was opposite of WT CD37 (Fig 2J, 2K). Taken together, CD37 that lacks glycosylation on position N183 is retained in the ER and therefore shows a lower surface expression.

### Glycosylation of CD53 does not affect its surface expression or nanoscale organization

Next, we investigated glycosylation of tetraspanin CD53 by mutating the glycosylation sites in a plasmid containing GFP-CD53. First, we verified that the surface expression of these constructs after transfection in CD53KO BJAB cells was similar to endogenous CD53 surface levels in BJAB cells (Fig 3A). Different from CD37, neither mutation of a single glycosylation site (N129Q, N148Q), nor mutation of both of them (NG) affected the surface expression of CD53 as detected by flow cytometry (Fig 3B, 3C). In line with these results, the non-glycosylated band that was present after transfection of WT GFP-CD53 as well as GFP-CD53(NG) could be detected by streptavidin after biotinylation-IP (Fig 3D, 3E) confirming their expression at the plasma membrane. These results were confirmed in experiments with ALFA-tagged CD53 (Fig S3A-D). The MW of the band pattern of the single mutants was between the fully glycosylated version of the WT (∼60 kDa) and the MW of non-glycosylated GFP-CD53 (∼45 kDa) (Fig S3E), suggesting that both glycosylation sites contribute to the full glycosylation of CD53. Although CD53 surface expression was not affected by its glycosylation, its turnover rate may still be altered. However, this possibility was excluded by internalization assays showing no glycosylation-dependent differences in CD53 internalization (Fig S3F, S3G).

**Figure 3:**
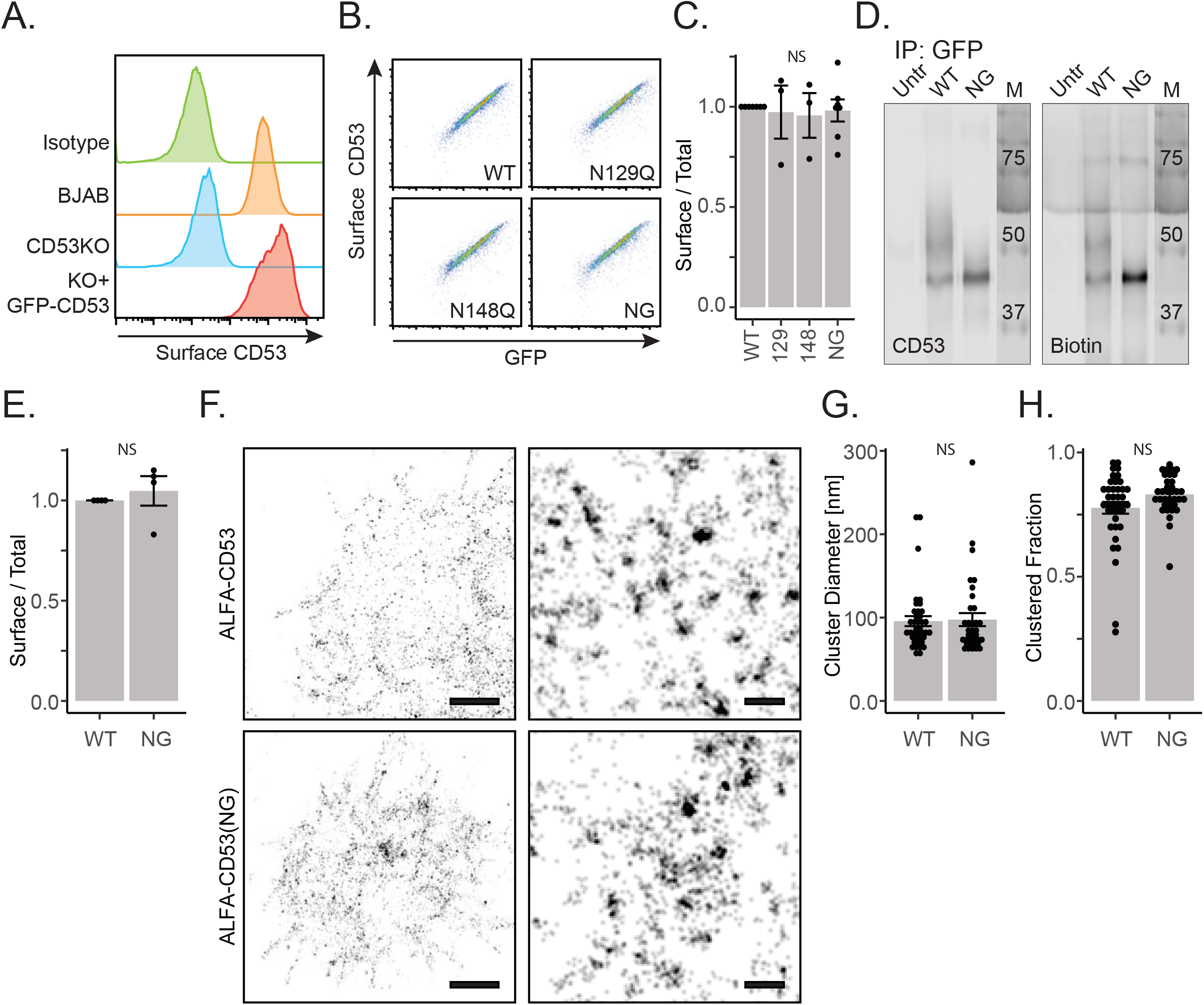
Glycosylation of CD53 does not affect its plasma membrane expression or nanoscale organization. (**A**) Endogenous surface expression of CD53 in BJAB cells compared to CD53KO BJAB cells transfected with GFP-CD53 analyzed by flow cytometry. (**B**) Correlation between antibody staining of surface CD53 versus GFP signal of CD53KO BJAB cells transfected with wild type (WT), single-mutant (N129Q, N148Q) or non-glycosylated (NG) GFP-CD53. (**C**) Quantification of (B). Antibody signal (surface) divided by GFP signal (total) normalized to WT (mean ± sem, Friedman test and Dunn’s multiple comparison, N=3-7). (**D**) Immunoprecipitation of WT and NG GFP-CD53 after surface biotinylation showing biotin signal in all CD53 bands. (**E**) Quantification of (D). Biotin signal (surface) divided by CD53 signal (total) normalized to WT (mean ± sem, Friedman test and Dunn’s multiple comparison, N=3). (**F**) dSTORM super-resolution microscopy images of the basal membranes of CD53KO cells overexpressing ALFA-CD53 (upper) or ALFA-CD53(NG) (lower). Right panels show zooms of the nanoscale organization of CD53. Scale bars: 2 μm (left) and 100 nm (right). Mean cluster diameter (**G**) and fraction of proteins residing in a cluster (**H**) was calculated per cell (black dots) based on Density-Based Spatial Clustering of Application with Noise (DBSCAN) analysis (mean ± sem, Friedman test and Dunn’s multiple comparison, N=5).

To assess the effect of glycosylation on the nanoscale organization of CD53, super-resolution dSTORM microscopy was employed to image the basal membrane of cells expressing either WT or NG ALFA-CD53 (Fig 3F). Both WT and NG CD53 were observed as a mixture of single as well as clustered proteins in the basal membrane, the latter being tetraspanin-enriched nanodomains. Average cluster size was analyzed by Density-Based Spatial Clustering of Application with Noise (DBSCAN) analysis as well as Pair-Correlation Analysis (PCA). CD53 was found in clusters with an average diameter of about 100 nm (Fig 3G, S3H) which is in agreement with Zuidscherwoude *et al* 2015 [27]. Loss of glycosylation did not affect the size of these clusters and neither the fraction of proteins residing in clusters (Fig 3H). Taken together, glycosylation does not influence surface expression and nanoscale organization of CD53 on the plasma membrane.

### Glycosylation differentially affects protein interactions of CD37 and CD53

Next, the effect of tetraspanin glycosylation on the interaction with known partner proteins was investigated. The interaction of CD37 with CD20 was not affected by mutating glycosylation sides N170 and N188 as shown by co-immunoprecipitation experiments (Fig. 4A). In contrast, mutation of N183 resulted in a significant decrease in CD20-CD37 interaction (Fig 4A, B) which was proportional to its decreased surface expression (Fig. 2E,F). Next, the interaction between CD37 and IL-6Ra was studied with the different glycosylation mutants. In contrast to CD37 interaction with CD20, no glycosylation dependency was observed for the interaction between CD37 and IL-6Rα (Fig 4C, D).

**Figure 4:**
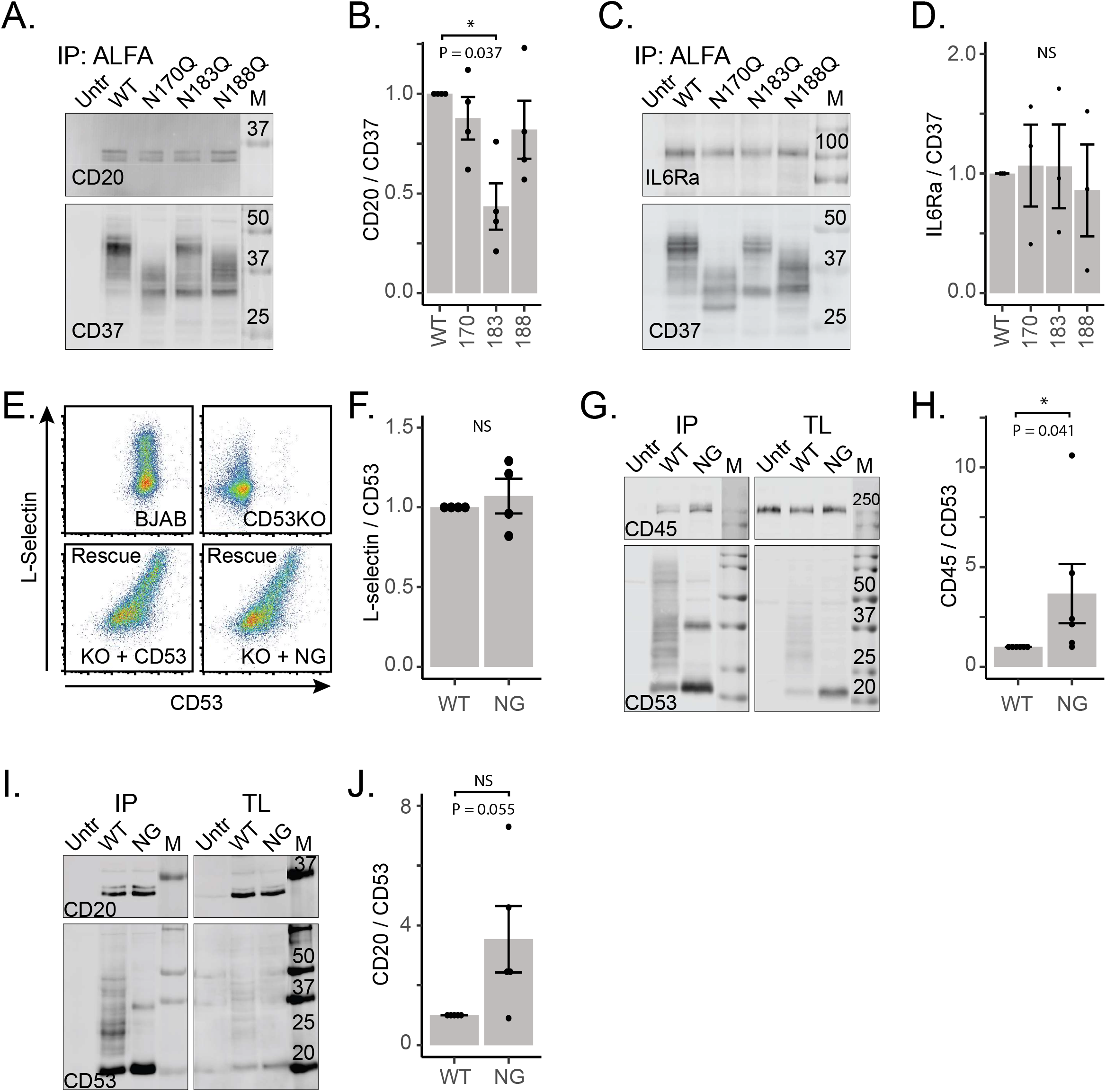
Glycosylation differentially affects protein interactions of CD37 and CD53. Immunoprecipitation (IP) of WT and single-mutant ALFA-CD37 expressed in CD37KO BJAB cells and co-IP of endogenous CD20 (**A**) or transfected IL-6Rα-GFP (**C**). Signal of co-immunoprecipitated CD20 or IL-6Rα-GFP divided by signal of immunoprecipitated CD37 normalized to WT (mean ± sem, Friedman test and Dunn’s multiple comparison, N=4) (**B**,**D**). (**E**) Flow cytometry analysis of surface overexpressed L-selectin, which is higher in BJAB cells compared to CD53KO BJAB cells. Co-overexpression of both WT and NG CD53 rescues L-selectin surface levels. (**F**) Quantification of (E). Surface L-selectin signal divided by surface CD53 signal normalized to WT (mean ± sem, non-paired T-test, N=4). Immunoprecipitation (IP) of WT and NG ALFA-CD53 expressed in CD53KO BJAB cells and co-IP of endogenous CD45 (**G**) or endogenous CD20 (**I**). TL=total lysate. Signal of co-immunoprecipitated CD45 or CD20 divided by signal of immunoprecipitated CD53 normalized to WT (mean ± sem, non-paired T-test, N=5) (**H**,**J**). * P≤0.05

Murine CD53-deficient lymphocytes were shown to be defective in lymph node homing as a result of loss of surface L-selectin [22]. In line with this study, transfected L-selectin could only be detected at the cell surface of CD53-expressing BJAB cells, and not CD53 KO cells (Fig 4E). Co-expression of either WT or NG CD53 rescued surface L-selectin expression to a similar extend, indicating that glycosylation is not involved in stabilizing L-selectin on the membrane (Fig 4E, 4F).

We recently discovered that CD53 interacts with CD45, a pan leucocyte marker involved in BCR and TCR signaling [24]. Co-IP experiments confirmed this interaction in BJAB cells, and surprisingly, loss of glycosylation enhanced CD53 interaction with CD45 (Fig 4G, H). To study whether this was specific for CD45 or also evident for other CD53 interaction partners, co-IP experiments were extended to CD20 [28]. Again, non-glycosylated CD53 was observed to interact better with CD20, although not significant (Figure 4I, F). Together these data show that glycosylation of CD53 inhibits the interaction with its partner proteins CD45 and CD20.

## Conclusions and Discussion

Many important cellular processes are modulated by tetraspanin proteins [1,2,29,30]. Most tetraspanins are highly glycosylated and this post-translational modification has been reported to affect their function in modulating the malignancy of several cancer types [10–12]. However, the function of tetraspanin glycosylation in immune cells remains largely unstudied. CD37 and CD53 are highly glycosylated tetraspanins that are both important in cellular and humoral immunity [14,21,31]. In the current study we found broad and cell-specific repertoires of glycosylated CD37 and CD53 in B cells. In line with our study, similar cell-specific glycosylation repertoires have been found for CD82 [32], Tspan3 [33] and CD63 [34]. In this study, we show how glycosylation differentially affects localization and function of CD37 and CD53 in B cells.

Whereas CD37 surface expression was clearly dependent on its glycosylation, this was not the case for CD53. The low surface expression of glycosylation-deficient CD37 and the lack of detection by EC2-directed antibodies suggests that glycosylation of CD37 is important for its structural integrity. It is possible that the glycosylation-dependent chaperones calnexin and calreticulin [35] are important for proper folding and localization of CD37. In contrast, no glycosylation dependency was observed for CD53 localization in line with studies on glycosylation of tetraspanin UPIb [36], CD63 [9,37], RDS [38] and CD82 [13]. The glycosylation-independent localization of CD82 is remarkable as we here report a clear glycosylation dependency for its closest homologue CD37. As far as we are aware only two other tetraspanins, Tspan1 and Tspan8, require glycosylation to obtain their normal localization [39,40].

Furthermore, we observed glycosylation to differentially affect interaction of CD37 and CD53 with their partner proteins. Whereas glycosylation did not affect CD37 interaction with CD20 and IL-6Rα, other than via changing subcellular localization, glycosylation was found to inhibit the interaction between CD53 and its partner proteins CD20 and CD45. Since cell surface levels of L-selectin were not affected by CD53 glycosylation, this inhibitory effect appears to be specific to certain partner proteins. We hypothesize that glycosylation sterically hinders the binding of partner proteins to specific epitopes on CD53. However, the flexible nature of the glycosylation groups may allow for them being pushed aside when the interaction is strong enough. Glycosylation may thereby select for strong, functional interactions. To our knowledge, inhibition of tetraspanin-partner interactions by glycosylation has not been reported before as other studies either found no effect, like RDS-ROM1 [38], or increased binding upon glycosylation, like CD63-CXCR4 [9] and CD82-integrin α5β1 [11].

Although it is unclear whether the observed inhibitory or stimulating functions of glycosylation are specific for the tetraspanin-partner interaction or intrinsic to the tetraspanin, it is tempting to speculate that different partner proteins favor specific sub-populations of tetraspanin proteins. This hypothesis would imply that the total pool of tetraspanin proteins in a cell is separated in many different functional groups based on the attached glycosylation groups. In line with this hypothesis, CD151 was shown to exclusively co-IP highly glycosylated CD82 [41], and MHC-I was found to interact exclusively with lowly glycosylated Tspan5 [42].

Our observation that glycosylation differentially affects the function of CD37 and CD53 is consistent with the few other reports showing tetraspanin-specific effects of glycosylation. In this limited set of studies, no clear correlation could be observed between the EC2 topology, the position of the N-glycosylation sites in the EC2 and the effect of glycosylation. For example, both CD37 and Tspan1 require glycosylation for their exit from the ER but they belong to different EC2 topology groups [43]. Moreover, although the position of glycosylation sites is rather similar for CD53 vs CD63 (both flanking the B helix in their EC2), and CD37 vs RDS (between the C and D helices of their EC2), the effect of glycosylation on localization and interactions is different.

Given the important role of CD37 in the onset and malignancy of B cell lymphoma [16–19], it is interesting to note that the glycosylation repertoires of CD37 in (malignant) B cell lines are broader than those observed in primary B cells and also reach higher molecular weights. Changes in CD37 glycosylation patterns could therefore be correlated to the onset or progression of cancer, as has been shown for CD82 [11,32]. Taken together, we report broad and cell-specific repertoires of glycosylated CD37 and CD53 in B cells, which adds another layer of regulation to tetraspanin-mediated membrane organization.

## Acknowledgements

We acknowledge funding support from the Netherlands Organization for Scientific Research (NWO): the Institute of Chemical Immunology (project ICI 000-23), ZonMW (project 09120012010023), the Dutch Cancer Society (project 12949 and 14726), and the European Research Council: Consolidator Grant (project 724281) and Proof-of-Concept Grant (project 101112687).

## Supplemental Figure legends

**Figure S1:**
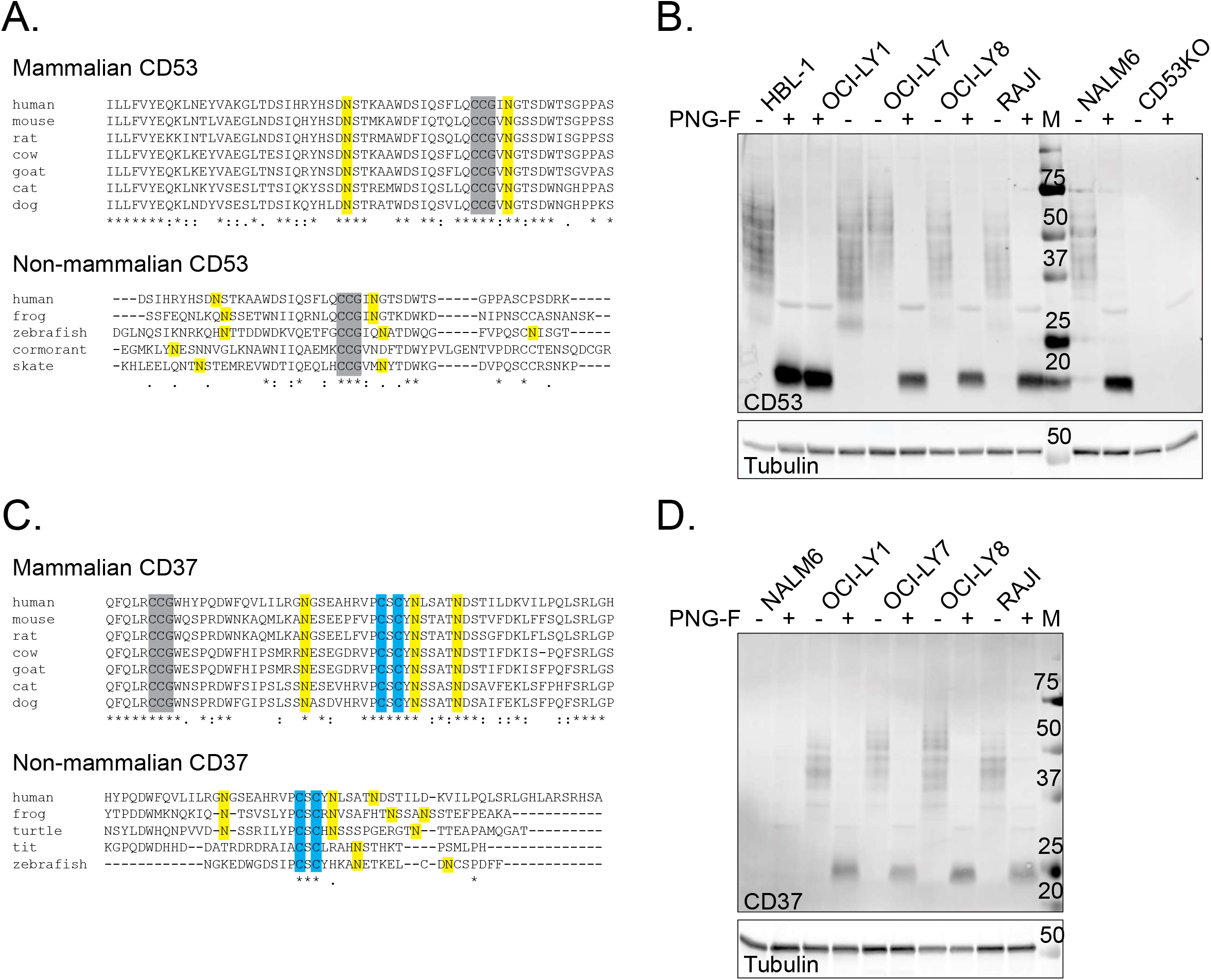
Evolutionary conservation and cell-specific usage of N-glycosylation sites in CD53 and CD37 in B cell lines. Multiple sequence alignment of both mammalian and non-mammalian CD53 (**A**) and CD37 (**C**) sequences revealing the high conservation of the N-glycosylation sites (in yellow). The tetraspanin-specific CCG-motif is highlighted in grey and additional cysteines in blue. Glycosylation of CD53 (**B**) and CD37 (**D**) is visible on western blot as a broad band pattern (25-70 kDa) in different B cell lines. CD53 knock-out NALM6 cells serve as negative control for the CD53 staining and WT NALM6 cells (a pre-B cell line) do not express CD37. Treatment with PNGase-F removes all N-glycosylation leading to a single band around 20 kDa (CD53) and 25 kDa (CD37).

**Figure S2:**
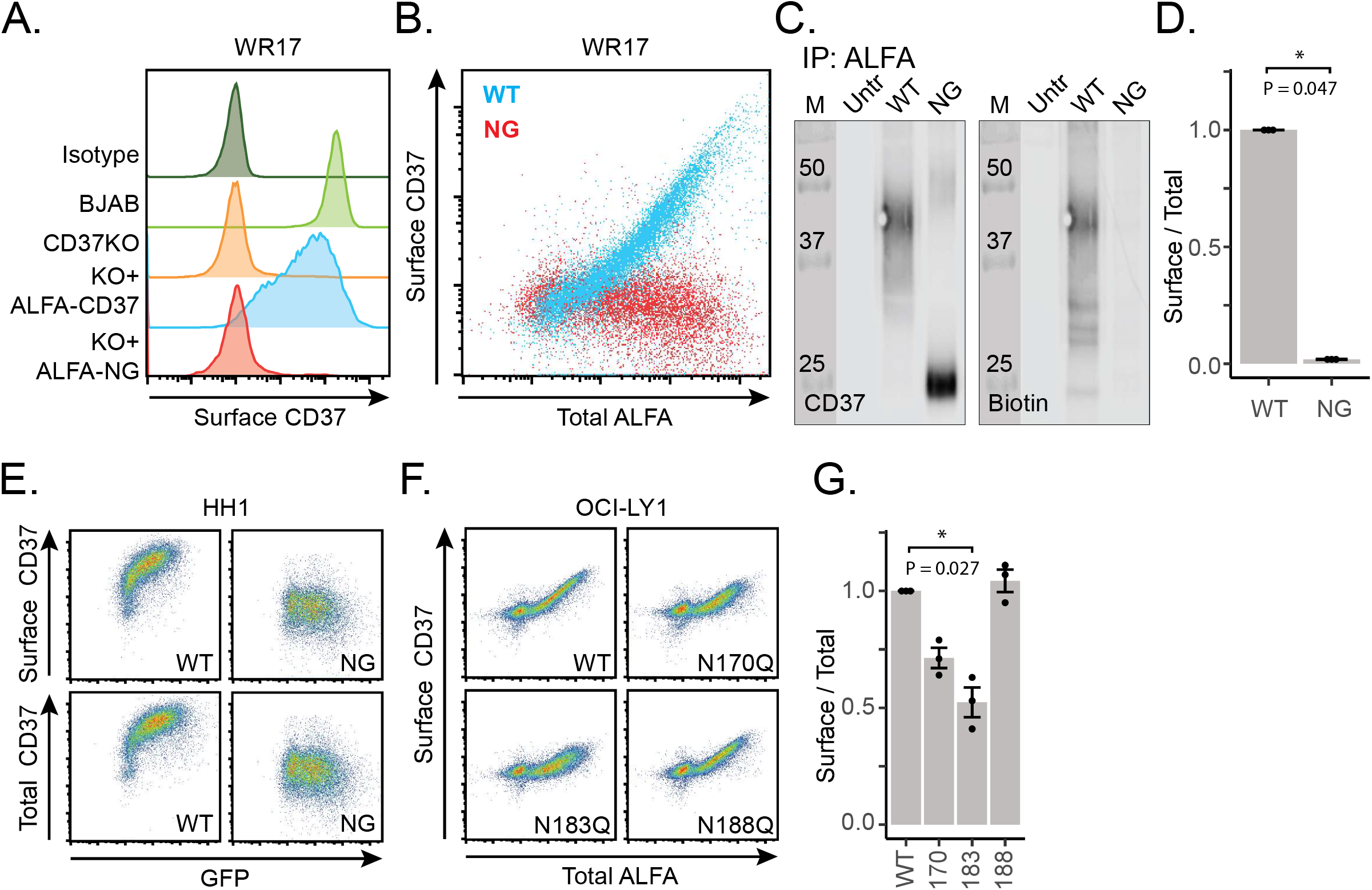
Surface expression of ALFA-tagged CD37 is dependent on its glycosylation. (**A**) Flow cytometry (based on the WR17 clone) showing endogenous surface expression of CD37 in BJAB cells compared to CD37KO BJAB cells transfected with ALFA-CD37 (WT) or its non-glycosylated version (NG). (**B**) Correlation between surface CD37 (antibody staining with WR17) and total CD37 (intracellular ALFA staining) of CD37KO BJABs transfected with WT or NG ALFA-CD37. (**C**) Correlation between antibody signal (HH1 clone) of surface and total CD37 versus ALFA signal of CD37KO BJABs transfected with WT or NG ALFA-CD37. (**D**) Immunoprecipitation of WT and NG ALFA-CD37 expressed in CD37KO BJAB cells after surface biotinylation. (**E**) Biotin signal (surface) divided by CD37 signal (total) normalized to WT (mean ± sem, non-paired T-test, N=3). (**F**) Correlation between surface CD37 signal (WR17 clone) versus total CD37 (intracellular ALFA signal) of CD37KO OCI-LY1 cells transfected with ALFA-CD37 or its single-mutant versions N170Q, N183Q and N188Q. (**G**) Antibody signal (surface) divided by ALFA signal (total) normalized to WT (mean ± sem, Friedman test and Dunn’s multiple comparison, N=3). * P≤0.05

**Figure S3:**
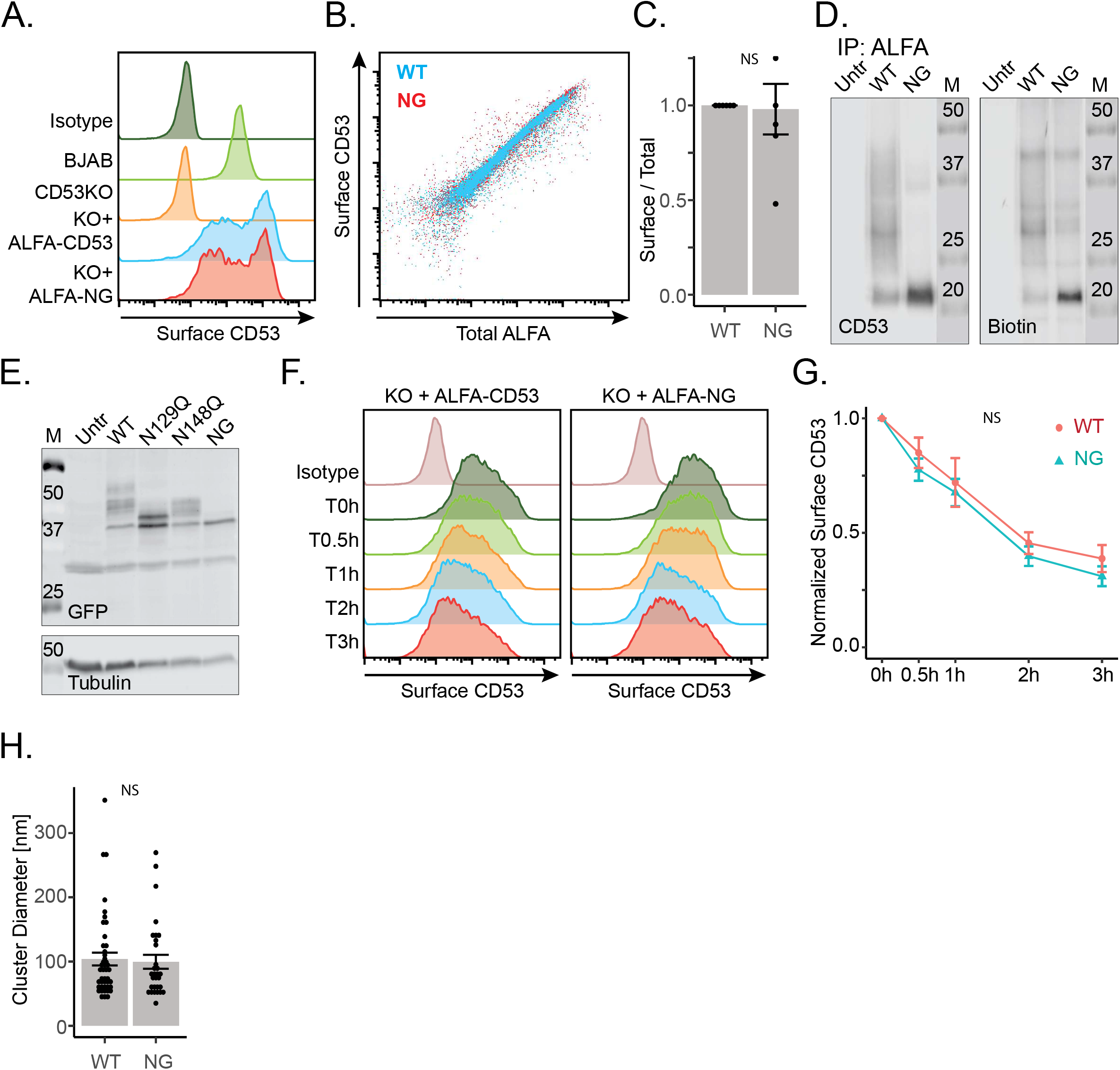
Surface expression and internalization of ALFA-tagged CD53 is glycosylation-independent. (**A**) Endogenous surface expression of CD53 in BJAB cells compared to CD53KO BJAB cells transfected with ALFA-CD53 (WT) or its non-glycosylated version (NG) by flow cytometry. (**B**) Correlation between surface CD53 and total CD53 (intracellular ALFA staining) of CD53KO BJABs transfected with WT (in blue) or NG (in red) ALFA-CD53. (**C**) Immunoprecipitation of WT and NG ALFA-CD53 expressed in CD53KO BJAB cells after surface biotinylation. (**D**) Biotin signal (surface) divided by CD53 signal (total) normalized to WT (mean ± sem, non-paired T-test, N=5). (**E**) Western blot showing WT, single mutant (N129Q, N148Q) and NG GFP-CD53 expressed in CD53KO BJAB cells. Tubulin was used as a loading control. (**F**) Flow cytometry analysis of the internalization of surface WT and NG ALFA-CD53 expressed in CD53KO BJAB cells. (**G**) Quantification of (F), CD53 surface signal normalized to T0 of WT (red) and NG (green) ALFA-CD53 (mean ± sem, N=3). (**H**) Pair correlation analysis of dSTORM microscopy data revealing the dominant cluster diameter per cell of WT and NG ALFA-CD53 (mean ± sem, non-paired T-test, N=5).

## Materials and Methods

### Culture and transfection of cell lines

The different B cell lines mentioned in this study were cultured in RPMI-1640 containing 10% fetal bovine serum, 1% antibiotics/antimycotics (AA), and 1% Ultraglutamine and kept at 37°C, 5% CO2. Five million BJAB or OCI-LY1 cells were transfected with 2 μg plasmid DNA using the SF Cell Line 4D-Nucleofector X Kit L (Lonza) and the AMAXA Nucleofector biosystem (BJAB: Program DS104, OCI-Ly1: program DN-100). After transfection, cells were seeded in growth medium without AA and incubated for 16-24 hours before further experiments were conducted. CD37- and CD53-knockout BJAB and OCI-LY1 cells were generated by CRISPR/Cas9 technology as described [44].

### Isolation of human primary B cells

Human primary B cells were isolated from buffy coats obtained from healthy volunteers. Pan B cells were isolated from peripheral blood leukocytes using the Pan B cell isolation kit according to manufacturer’s instructions (Miltenyi Biotec).

### Flow cytometry

Cell suspensions containing 2×10^5^ cells were blocked for 30 minutes on ice in PBS containing 1% BSA, and 2% human serum. Incubations with primary antibodies were then performed in blocking buffer for 30 minutes on ice using the antibodies and dilutions indicated in Table S1. After two washes with blocking buffer incubations with fluorescently labeled secondary antibodies were then performed in blocking buffer for 30 minutes on ice using the antibodies and dilutions indicated in Table S1. After another two washes in PBS containing 1% BSA, samples were analyzed on a Cyan flow cytometer (Beckman Coulter) or a BD FACSLyric Flow cytometer (BD Biosciences), and data were analyzed using FlowJo X software (Tree Start Inc.). In case of intracellular staining, cells were fixed by incubation in 4% paraformaldehyde (PFA) in PBS for 30 minutes on ice and all subsequent buffers for blocking, staining and washing contained 0.5% saponin.

### Immunoprecipitation, enzyme treatment and Western blot

For immunoprecipitation, 5×10^6^ transfected cells were lysed in 1 ml 1% Brij97, 10 mM TRIS pH 7.5 and 150 mM NaCl supplemented with either 1 mM EDTA (for the CD53-CD45 co-IP’s) or 2 mM CaCl_2_ + 2 mM MgCl_2_ (for all other IP’s). After 30 min incubation on ice with intermittent vortexing, insoluble material was removed by centrifugation at 10,000 × g and the cell lysate was pre- cleared by addition of heat inactivated goat serum and protein G sepharose beads (GE Healthcare). Proteins were then either immunoprecipitated by adding 2 μg mouse anti-GFP (Clones 7.1 and 13.1, Roche) and protein G sepharose beads or ALFA selector ST beads (NanoTag Biotechnologies). In some cases immunoprecipitated proteins were treated with PNGase-F or Endo-H (New England Biolabs) according to the manufacturers protocols. Reduced immunoprecipitated proteins were separated on a 10 or 12% SDS-polyacrylamide gel and transferred to a PVDF membrane (GE Healthcare). Details on the blocking of the membranes and detection of the proteins can be found in Table S1. Acquisition was performed using either the Odyssey Infrared Imaging System (LI-COR Biosciences) or the Typhoon 5 (Amersham). Quantifications were performed using FIJI.

### Confocal microscopy and image analysis

Half a million transfected cells were adhered to poly-L-lysine-coated coverslides and subsequently fixed by incubation with 4% PFA in PBS for 30 minutes. Cells were blocked in 3% BSA, 1% filtered human serum, 10 mM glycine, and 0.5% saponin in PBS for 1 h. Cells were subsequently stained for 30 min in the same solution supplemented with 10 μg/ml primary antibody, and after thorough washing an appropriate secondary antibody (Table S1). Samples were then stained with DAPI, washed, and embedded in mowiol. Imaging was performed on an Olympus FV1000 Confocal Laser Scanning Microscope and quantification of co-localization was performed using the JACop-pluging of FIJI.

### dSTORM microscopy and data analysis

dSTORM microscopy samples were prepared by adhering 5×10^6^ cells to poly-L-lysine-coated coverslides and fixation with 0.1% glutaraldehyde, 4% PFA in 0.2 M phosphate buffer pH 7.4 for 1h. After thorough washing with PBS, remaining reactivity of the fixatives was quenched by a 30-min incubation with 100 mM glycine, 100 mM NH_4_Cl and 0.1% triton-X100 in PBS. Samples were then blocked for 1h in filtered 10 mM glycin, 10% NGS and 0.1% Triton-X100 in PBS and stained for 1h in filtered 10 mM glycin, 3% NGS and 0.1% Triton-X100 containing A647-tagged FluoTag-X2 anti-ALFA nanobody (NanoTag Biotechnologies). After extensive washing with both staining solution and PBS, samples were fixed by incubation in 4% PFA in PBS for 30 minutes and stored in 0.1% PFA in the fridge. Before imaging, samples were washed in PBS, quenched with 100 mM glycine in PBS for 15 minutes and again washed in PBS. Coverslips were then mounted in a custom-made low-drift magnetic sample holder and cells were imaged in 1 ml of OxEA buffer [45]. dSTORM microscopy was performed on a custom-build low-drift inverted microscope setup as described in Neviani *et al* [46].

Data sets were analyzed with FIJI IMAGE J 1.52g and the analysis module THUNDERSTORM [47]. A detection threshold of 300 photons was used, and typical uncertainty mode values for the localizations were 20 nm. Images were reconstructed using the averaged shifted histograms method with a rendering pixel size of 10 nm. The output of THUNDERSTORM, a data file containing the filtered localizations, was further analyzed by pair correlation analysis and the DBSCAN cluster algorithm (ε-value of 50 nm and a minpts of 5) in R.

### Plasmids and mutagenesis

Expression constructs containing wild type CD53 and CD37 with a GFP-tag fused to their C-terminus in the psGFP2-C1 vector have been described before [18,23] and were used as a template for further cloning. Single, double and triple N>G mutations were introduced in the parental constructs by using the Q5 Site-Directed Mutagenesis Kit (New England Biolabs) according to the manufacturers protocols and using custom designed primer. ALFA-tagged derivatives were created by a two-step cloning procedure. First, the CD53 or CD37 sequence was cut out of pSGFP2-C1 using EcoRI-HF and SalI-HF and ligated in pCDNA3.1(+) that was cut using EcoRI-HF and XhoI-HF (all enzymes used from NEB). Than site-directed mutagenesis was performed to introduce the ALFA tag as described in [26]. All constructs were verified by Sanger sequencing.

### Statistics

Bars and error bars represent the mean and standard error of the mean throughout the manuscript. Statistical testing was always performed on the non-normalized data and is indicated in the figure legends. Statistical significance was defined as * P≤0.05, **P≤0.01 and ***P≤0.001.

